# Simulations on the efficacy of radiotherapy with different time schemes of antiangiogenic therapy

**DOI:** 10.1101/2021.09.06.459137

**Authors:** Mert Tuzer, Defne Yilmaz, Mehmet Burcin Unlu

**Affiliations:** Department of Physics, Bogazici University, Bebek, Istanbul, Turkey; Center for Life Sciences and Technologies, Bogazici University, Bebek, Istanbul, Turkey; Global Station for Quantum Medical Science and Engineering, Global Institution for Collaborative Research and Education (GI-CoRE), Hokkaido University, Bebek, Sapporo 060-8648, Japan

## Abstract

The combination of radiotherapy and antiangiogenic agents has been suggested to be potent in tumor growth control compared to the application of antiangiogenic therapy or radiotherapy alone. Since radiotherapy is highly dependent on the oxygen level of the tumor area, antiangiogenic agents are utilized for the reoxygenation of tumor vasculature. We present a mathematical framework to investigate the efficacy of radiotherapy combined with antiangiogenic treatment. The framework consists of tumor cells, vasculature, and oxygenation levels evolving with time to mimic a tumor microenvironment. Non-linear partial differential equations (PDEs) are employed to simulate each component of the framework. Different treatment schemes are investigated to see the changes in tumor growth and oxygenation. To test combination schedules, radiation monotherapy, neoadjuvant, adjuvant, and concurrent cases are simulated. The efficiency of each therapy scheme on tumor growth control, the changes in tumor cell density, and oxygen levels shared by tumor cells are represented. The simulation results indicate that the application of radiotherapy after antiangiogenic treatment is more efficient in tumor growth control compared to other therapy schemes. The present study gives an insight into the possible interaction and timing of the combination of radiotherapy and antiangiogenic drug treatment.

## Introduction

Tumor microenvironment, a temporal and spatial complex system, is a major challenging factor in cancer treatment. In this environment, tumor blood vessels, which differ from normal vasculature in terms of organization, function, and hierarchy, play a significant role in cancer evolution^1^. The disorganized architecture of tumor vascular system leads to the heterogeneous blood flow and perfusion. This heterogeneous perfusion results in disruptions in the diffusive transport of oxygen and other nutrients into the tumor center, yielding hypoxic and acidic regions that trigger the secretion of growth factor inducing angiogenesis, which is a process for tumor cells to form a vascular network from the existing one for its rapid growth and migration.^2–5^.

To deal with the factors affecting the treatment response unfavorably in tumors, conventional cytotoxic therapies as radiotherapy and chemotherapy are applied to control the tumor growth. Strategies that aim to target the inhibition of tumor vasculature, known as antiangiogenic therapy, are also used to promote the outcome of conventional treatments. In addition, the combination of these therapies are employed in order to yield favorable results.

Radiotherapy is one of the treatment modalities being substantially benefited in cancer research^6^. Although conventionally applied to patients, its efficiency is most likely reduced by radioresistance, which might be due to internal factors such as genetic predisposition or external factors such as hypoxia. Therefore, it fails to cure some types of cancer when applied alone^7, 8^. Radiotherapy is commonly used with chemotherapeutic agents, but additive cytotoxicity to healthy tissues remains a serious problem^9, 10^. In order to overcome this problem, the combination of radiotherapy with different targeted therapies is investigated. Since oxygenation of a tumor region affects the outcome of treatment in favor of the radiation applied, hypoxia is a crucial factor in radioresistance that impedes the success of radiotherapy^11, 12^. As its name suggests, antiangiogenic agents inhibit angiogenesis, temporarily allowing abnormal tumor vessels to function in a normal manner. Considering this effect of agents on the vascular network, it might be counter-intuitive to think that a therapy benefiting antiangiogenic agents is also a candidate to combine with radiotherapy because of the aforementioned relationship of oxygenation and effectiveness of radiotherapy.

Studies have shown that the use of antiangiogenic agents can make endothelial cells sensitive to irradiation; therefore, administration of these agents during the time of irradiation enhances the cytotoxic effects of radiation therapy by leading to the endothelial cell killing induced by radiation and by improving tumor response^13, 14^. Moreover, antiangiogenic agents can improve the antitumor effect of radiation by inducing to reduce the vascular density and control the tumor growth delay^15, 16^.

In order to investigate the effects of the use of antiangiogenic agents together with radiotherapy, several experiments are conducted. In a human tumor xenograft of small cell lung carcinoma 54A, it is stated that the administration of DC101, the antibody of anti-vascular endothelial growth factor receptor-2 (anti-VEGF2), together with radiotherapy has a remarkable combined effect that results in an improved tumor growth control compared to the radiotherapy alone^17^. In another study led by Becker *et al*.^18^, bevacizumab, an antiangiogenic agent, is injected before and after radiotherapy in different experiments of a lung adenocarcinoma model in mice. The injection of antiangiogenic agent before two hours of radiation therapy has shown promising results on tumor response and an increase in necrotic regions. In a study led by Ansiaux *et al*.^19^, it is reported that Thalidomide, a type of antiangiogenic agent, can radiosensitize the tumor cells by promoting a reoxygenation period within two days.

Radiation therapy induces changes in the blood vessels depending on the dose and fraction size applied^20^. With the decreased vessel density, the distance between the functional vessels increases, causing reduced tissue perfusion. In addition, irradiation leads to an increase in the permeability and leakiness of tumor vessels and makes them more radiosensitive compared to the surrounding healthy tissue^21^. Normal vessels are highly radioresistant compared to immature vessels, i.e. newly forming ones, so tumor vasculature, which is mostly composed of these immature vessels, is more vulnerable to radiation^22^.

The effects of both single-dose and fractionated radiotherapy on tumor perfusion and oxygenation are based on dosing and scheduling^23–25^, making these variables crucial for the combination of radiotherapy with antiangiogenic agents. It is stated that the application time and dose rates of radiotherapy and antiangiogenic agents as well as their sequences (adjuvant, neoadjuvant, or concurrent) have a pivotal role in determining the effectiveness of the treatment^26^. In a study of breast cancer and ovarian carcinoma, it is shown that tumor growth is impeded in combination therapy but the increase in oxygen levels inside the tumor is only seen between days 2 and 5^27^. In a squamous cell carcinoma model, oxygenation is increased for several days compared to the control case, but decreased when the tumor is given sunitinib after exposure to a single dose of radiation^28^. In addition, several preclinical and clinical studies have stated that the maintenance administration of antiangiogenic agents during and after radiation therapy has the potential for the inhibition of tumor regrowth^29, 30^.

Although experimental and clinical studies are conducted to determine the feasible approach regarding dosages, scheduling, and sequencing to enhance the treatment results, mathematical models might be more convenient and applicable to optimize treatments. Various treatment scenarios with different combinations of therapies could be tested utilizing properly developed mathematical models before clinical trials. There are several studies on the mathematical modeling of these two monotherapies and the combined therapy seeking the optimal sequencing of therapies. In a non-spatial model, a two-compartmental system of tumor and vasculature is studied to temporally optimize treatments, which lacks the factor of oxygen and complex interactions between tumor and vasculature^31, 32^. In another study, a spatio-temporal model, including tumor cell density, vasculature, and concentration of oxygen and nutrients, has been constructed to find suitable dosages and treatment schedules^33^.

In this study, we build a framework that includes radiation therapy and antiangiogenic therapy, following the mathematical model composed by^34^. We use the model to investigate the effects of combination therapy on tumor regression. In the model, different schedules of radiation therapy and antiangiogenic therapy are also examined to explore the optimal sequencing of combination therapy.

## Results

In this study, we examine the interaction between radiation therapy and antiangiogenic agents under various schedules. Since it is important to find feasible treatment schedules of the combination therapy, different treatment regimes are investigated in an effort to see the changes in tumor growth and oxygenation.

For all schedules, a 2-D mathematical model is used and the model begins with a tumor whose density is distributed to be Gaussian. The initial radius of the tumor is about 0.2 mm, and after evolving 30 days, the tumor is left to grow up to a radius of about 10 mm, which is consistent with the benefited model^33^. Vessels and oxygen levels are randomly and positively distributed and both are dependent on tumor and each other, thus, displaying a heterogeneous picture in subsequent iterations.

In the model, vessels are distributed randomly in the initial state and become heterogeneous because of the tumor cells (see Figure 1, vessel density). With the increase in tumor cell density, vessels become sparser in the inner part of the tumor; however, angiogenesis causes increased vessel density. The density of tumor cells is highest in the tumor center. As the distance to the neighboring healthy tissue increases, the tumor cell density decreases. Moreover, due to the decreased oxygen concentration in the interior parts of the tumor, necrotic regions can be observed in these areas (see Figure 1, tumor density).

**Figure 1.**
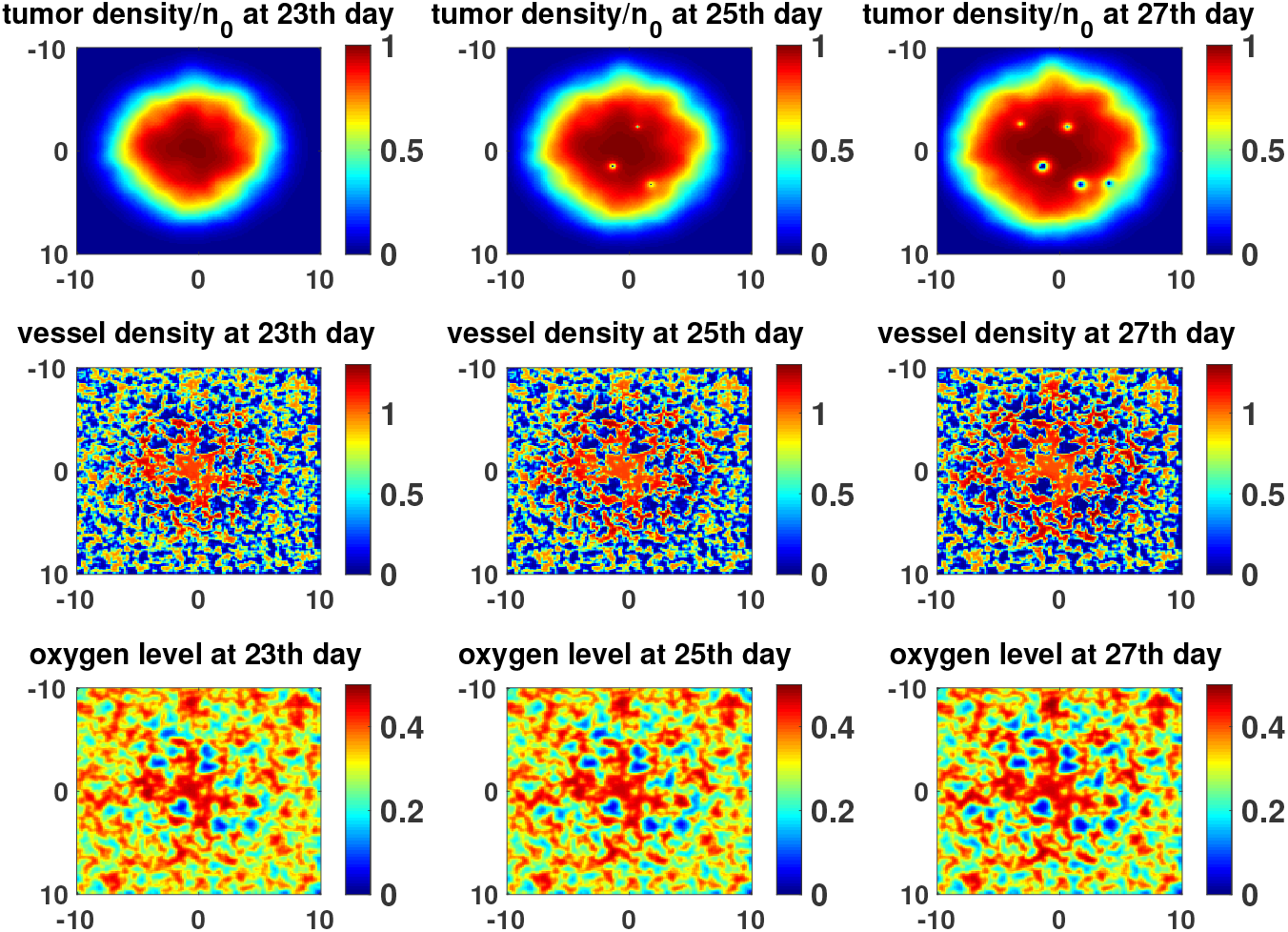
Tumor growth, vasculature and oxygen for control case. Tumor density, vessels and oxygen level are depicted for days 23, 25 and 27.

To test combination schedules, radiation monotherapy, neoadjuvant, adjuvant, and concurrent cases have been simulated, where radiotherapy alone is applied, antiangiogenic agents are injected before, after, and concurrently with radiotherapy, respectively. Since vessel values are initially randomly distributed, each treatment case has been simulated with ten different vessel networks and the calculated quantities (total tumor density and effective oxygen level) have been averaged. During simulations, the application of radiotherapy is always started from day 18 to day 20 in daily fractions where the tumor has a radius of about 1.5 mm, as in the case in the following experiment^35^ and the mathematical model^33^. Antiangiogenic treatment is also given in 3 daily pulses with five different timings starting from days 12, 15, 17, 21, and 24 to investigate neoadjuvant, concurrent, and adjuvant cases. By employing different treatment schedules, the distributions of tumor cells are represented in Figure 2. Each treatment schedule has been initialized with the same tumor and vascular network in order to monitor the effects more accurately. Compared to adjuvant cases, early injection of antiangiogenic agents (neoadjuvant cases) has shown more pronounced results in tumor control (see Fig 2), which can be easily observed with the change in necrotic regions.

**Figure 2.**
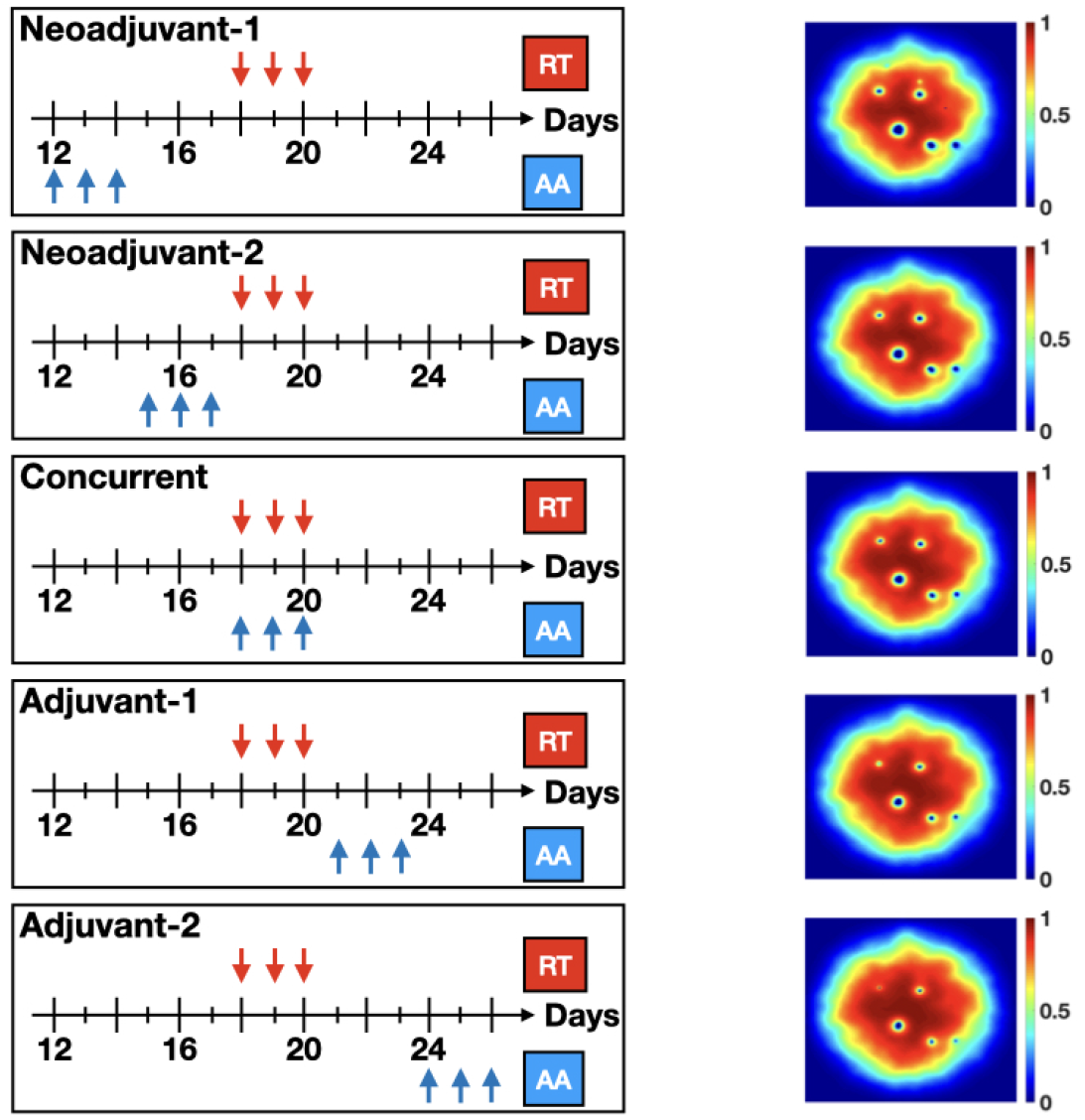
Treatment regimens applied and tumor density changes for these regimens. Tumor density for day 27 is calculated for different cases.

The efficiency of each therapy scheme on the tumor growth control is depicted in Figure 3, which shows the ratio of the daily total tumor cell density to the initial total tumor cell density, where the total tumor cell density for each day can be calculated as follows:

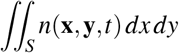

within the area S, bounded with the condition n(**x,y**,t)>0.1, covering the tumor region.

**Figure 3.**
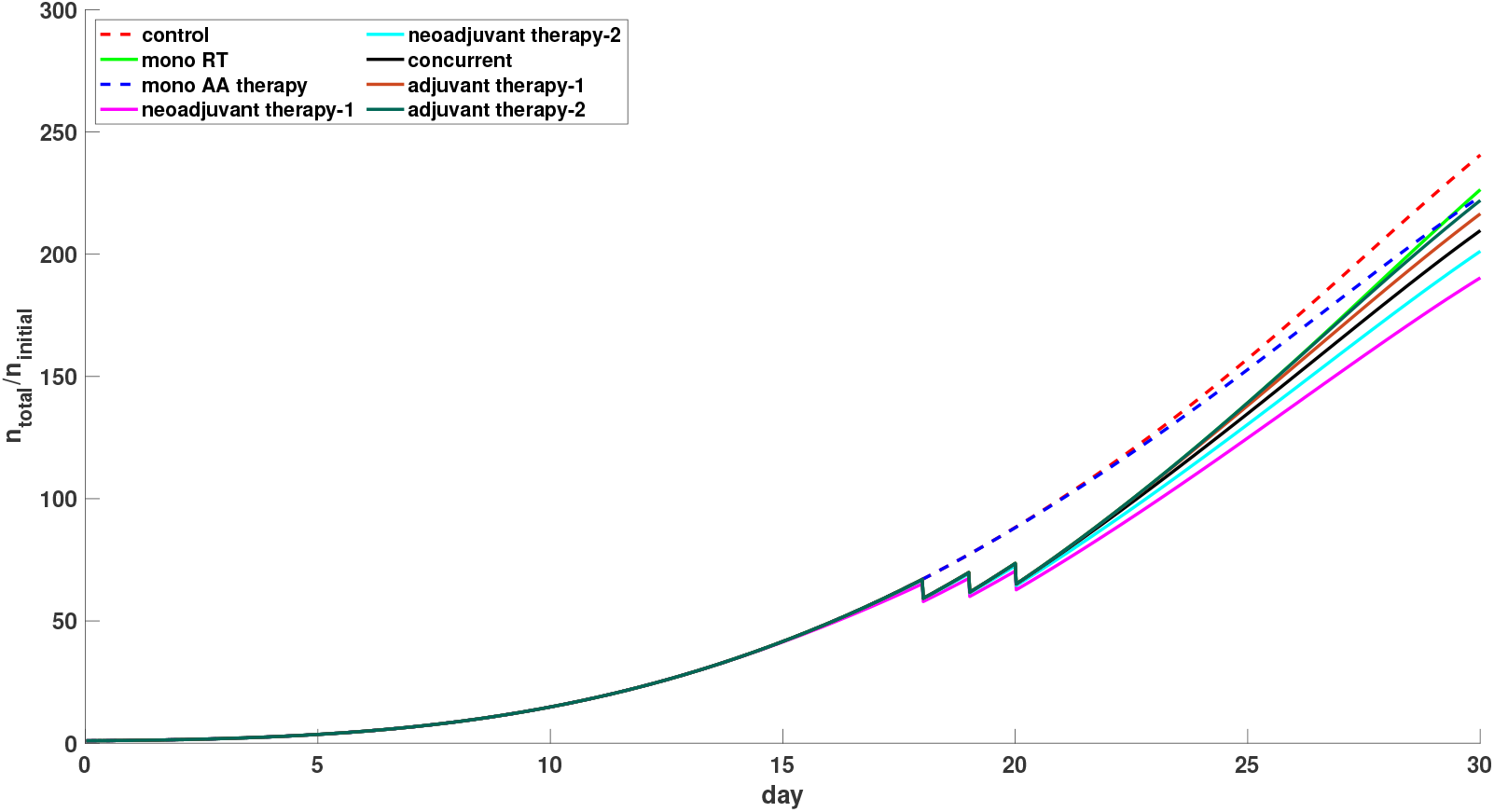
The daily total of tumor cells normalized with the one on the initial day.

The antiangiogenic agents affect the total tumor cell density, not at the first moment but after a few days, since the treatment directly acts upon the vascular system and oxygenation levels, and then the effect shows itself in the tumor cell density. As can be seen in Figure 3, total tumor cell density has started to decrease on day 23 in the case of antiangiogenic treatment alone. The cases of radiation monotherapy, adjuvant and concurrent therapies start with the same effect since radiotherapy is applied on the same day. Among the combined therapies, application of antiangiogenic agents before radiotherapy is more effective in tumor growth control compared to the other three cases; while the injection of antiangiogenic agents after radiotherapy has almost the same effect in growth control as the use of antiangiogenic agents or radiotherapy alone.

As it is expected, antiangiogenic treatment does not affect the tumor growth control profoundly when it is applied alone (see Figure 3). Since only the density of vasculature is used in this model, which lacks the complex interactions of vessels and antiangiogenic agents, the modeling of microvessel diameters and pore sizes should be provided to observe a transient reoxygenation period or the normalization window clearly, not existing in this mathematical framework. Hence, rather than alleviation of hypoxia, a decrease in oxygen concentration shared by tumor cells is observed after antiangiogenic agents are injected in the simulations.

It has been stated that low-dose radiation doses applied in the daily fractionation scheme in conventional radiation therapy induces a reoxygenation period^36^. To gain an insight on the oxygen levels shared by the tumor cells and to investigate the oxygen levels in the cases used in our model, a parameter called the *effective oxygen level* is introduced. This parameter can be defined as:

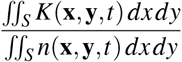

Following the equation above, Figure 4 is presented to examine the changes in the effective oxygen levels for each therapy. After every radiotherapy session, a small increase in oxygen levels is observed. Since antiangiogenic therapy is modeled to decrease vessel density, it directly reduces oxygen levels. This effect can be observed from Figure 4 if all other cases are compared with radiation monotherapy. Furthermore, the case of only antiangiogenic agent administration in the figure shows the reduction in effective oxygen levels after injections. At the administration of the earliest injection of antiangiogenic agents, the effective oxygen level has the lowest value, while all combination cases eventually approach the value of the control case.

**Figure 4.**
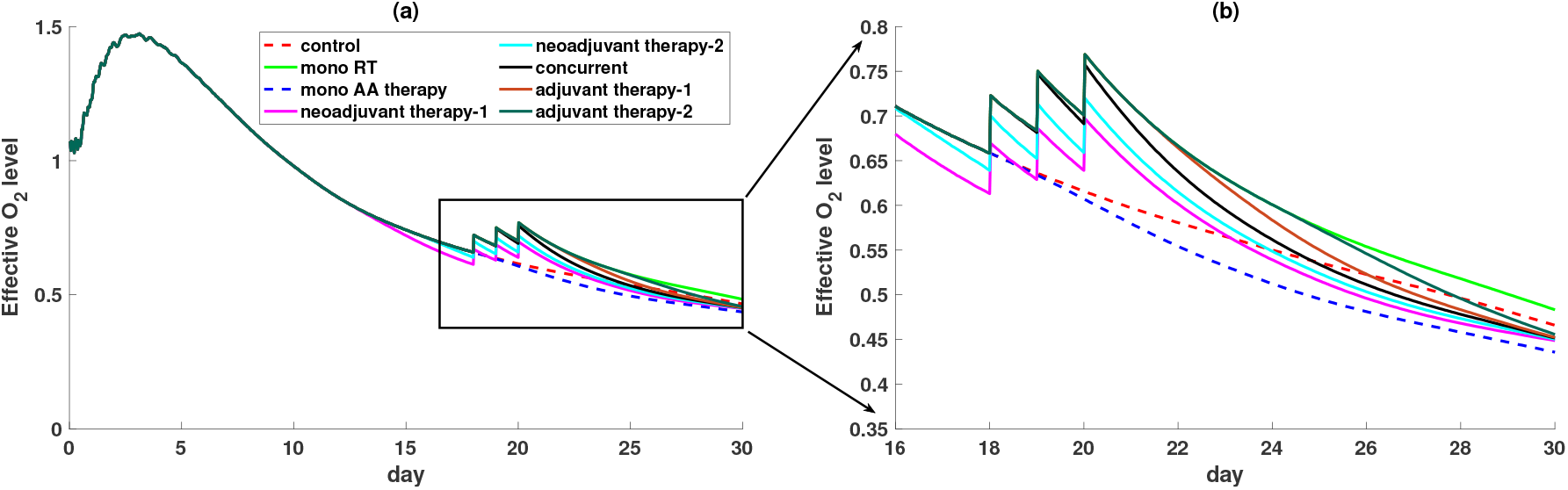
Effective oxygen levels shared by tumor cells. **(a)** The effective oxygen levels within days. **(b)** The zoomed version within the days of therapy sessions.

To investigate the dependency of selecting parameters such as tumor cells-radiotherapy reaction coefficient, *γ*_1_, and vasculature-antiangiogenic agent reaction coefficient, *ζ*_3_, the sensitivity of these parameters has been analyzed as shown in Table 1 and depicted in Figure 5. While the dependency of change in *γ*_1_ is explored in the case of radiotherapy alone, the change in *ζ*_3_ is studied in only antiangiogenic therapy case. In Table 1, N is a denotation for the total tumor density, and the regression is a comparison of how much tumor growth is retarded in days compared to the control case. As can be seen in Table 1 and Figure 5, the density of tumor cells decreases and the regression of tumor growth increases for radiotherapy alone case as the effect of radiotherapy on tumor cells increases. In the only antiangiogenic therapy case, the regression in tumor growth and the changes in tumor cell density are more pronounced with the increased reaction of antiangiogenic agents on the vasculature.

**Table 1.**
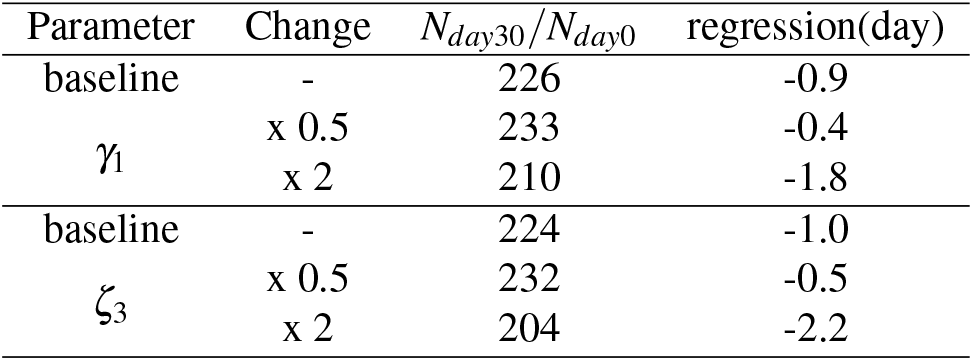
Parameter sensitivity analysis for *γ*_1_ and *ζ*_3_

**Figure 5.**
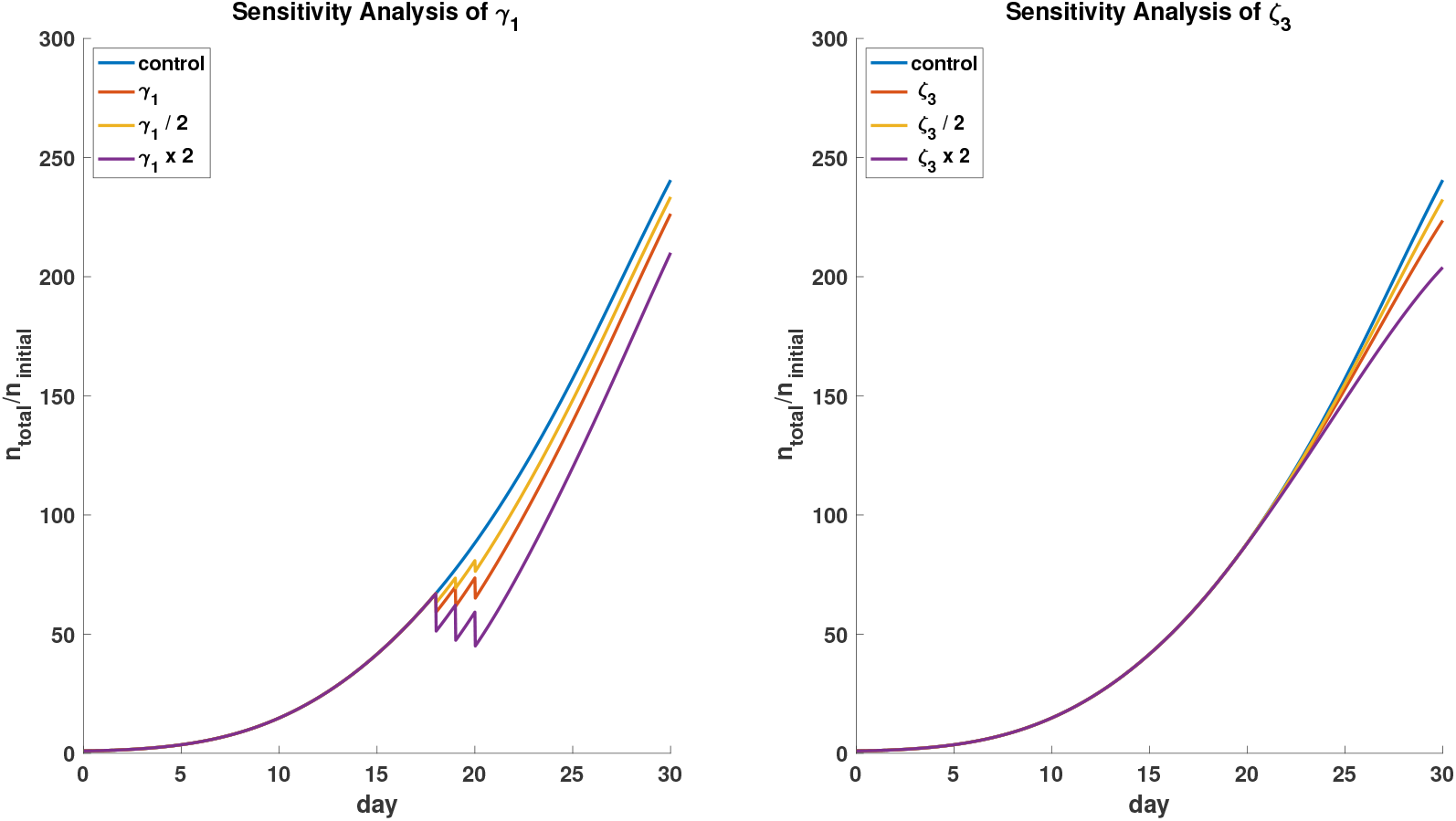
Sensitivity analyses of tumor cells-radiotherapy reaction coefficient, *γ*_1_ and vasculature-antiangiogenic drug reaction coefficient, *ζ*_3_.

## Discussion

We investigate the efficacy of the combination of antiangiogenic agents and radiotherapy with different treatment schedules, employing our mathematical framework. This framework consists of a tumor site, a vascular network, and oxygenation levels evolving with time to mimic a tumor microenvironment. As can be figured out from the model equations, these three layers are interdependent with each other. That is, oxygen is needed for the growth of a tumor, and vessels are used to feed oxygenation levels. On the other hand, the growing tumor invades the vascular system and annihilates with a predetermined factor, and consequently the oxygen levels within that region decrease in addition to the consumption of oxygen due to the tumor itself.

The relationship between radiation therapy and antiangiogenic therapy is investigated by several preclinical and clinical studies. In an experiment by Lee *et al*., it is found that tumor oxygenation increases when antiangiogenic agents are applied together with radiation therapy and this enhancement has been associated with the decrease in oxygen consumption due to the increased tumor cell death^15^. In our study, we also observe the same effect such that radiation therapy followed by antiangiogenic agents results in an improved tumor growth control and this result comes from an increase in necrotic regions thereby a decrease in oxygen levels following the application of the antiangiogenic agent.

In combination therapy, it is crucial to optimize the timing of treatments. In this context, several studies have emphasized the importance of appropriate scheduling of radiation therapy and antiangiogenic therapy^30, 37^, indicating that the tumor response varies to the combination therapy depending on the variable duration of administration of both treatments. Similar to our results, it is stated that co-administration of radiotherapy and antiangiogenic agents and radiotherapy before antiangiogenic agents are less effective in tumor control, however; when radiation therapy is applied after antiangiogenic agents, an enhancement is observed in the growth control. They have also suggested that the application of radiation after antiangiogenic therapy yields more effective results than radiation or antiangiogenic therapies alone^30^.

It has been documented before that bevacizumab, a commonly used antiangiogenic agent, given together with dose-intensive radiotherapy carries a risk of severe and unusual toxicity once irradiating the mucosal organs such as bronchi, intestines, and urethra^38–41^. The detrimental effect on the mucosa, mostly with hypofractionated high dose regimens, seemed to be true notwithstanding the order and time between the radiotherapy and bevacizumab. However, there are also other publications revealing the safety of bevacizumab and radiotherapy combinations^42, 43^. Aside from its antiangiogenic effect to improve tumor response, bevacizumab has been increasingly used to clinically control, stabilize, and hopefully revert the clinically symptomatic radiation-induced brain necrosis^44–47^. Therefore, it is currently unclear what is long enough to be a safe interval in a sequence of given bevacizumab to radiotherapy, and how the antiangiogenic effect can be paired with radiotherapy.

Our simulation results indicate that the neoadjuvant scheduling of the combination therapy is more efficient in tumor growth control compared to other schedules. When antiangiogenic agents are applied after radiotherapy, the total number of tumor cells is almost equal to the case in which radiotherapy is applied alone. Besides, administration of antiangiogenic agents alone has no effect on tumor growth control, as expected, and exhibits similar behavior to the control case.

In the model, only the vessel density is taken into account so the complex interplay between the vascular network and antiangiogenic agents is not considered. In the simulations, the application of antiangiogenic agents results in a decrease in oxygen concentration of tumor cells rather than an alleviation of hypoxia. However, a detailed structure including the pore sizes and the diameters of microvessels might be established as future work to determine the normalization window clearly.

## Methods

In this model, non-linear partial differential equations (PDEs) are employed for each component and the equations are solved by using the finite difference method. In the following equations, tumor cells, vessel density, and oxygen levels are denoted by n, m, and K respectively.

### Tumor cells and radiotherapy

The equation (1) is used to describe the spatio-temporal changes in tumor cell density. In the equation, the first term is the diffusion of tumor cells, where D_n_ is the diffusion coefficient of tumor cells and the second term is applied to describe the growth of tumor cell density up to a carrying capacity n_lim_, where *ρ* is the growth rate. In the case where only these two first terms are present, the equation (1) has two fixed points such that at n=0, there is an unstable fixed point in which there is not any cell population and at n=n_lim_, the cell population approaches its carrying capacity.

The third term in equation 1 is employed for the interaction of tumor cells between low-dose fractionated radiotherapy. Here, the parameter *h*_*n*_(**x**, *t*) represents a map for oxygen enhancement ratio (OER) dependency, which is known to be in a range of 2-4 for low-LET radiations such as X-rays. For this purpose, oxygen levels are simply mapped to the range of 1 to 3 linearly. The last term, *γ*_2_*n*(**x**, *t*)*K*(**x**, *t*), accounts for the interaction between tumor cells and oxygen in blood vessels. Tumor cells proliferate at a rate *γ*_2_ when they are supplied with oxygen.

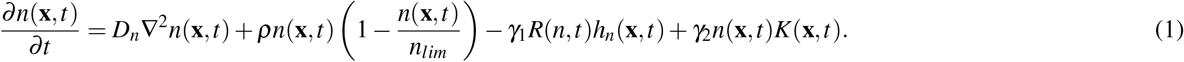

In the linear quadratic model, which is mostly used in radiotherapy, the decay rate *R*(*n, t*) can be defined as^48^:

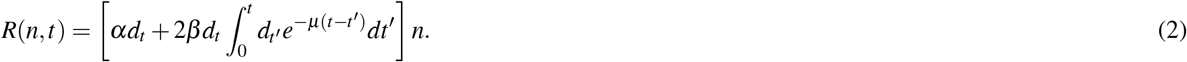

Here, *α* and *β* are the sensitivity parameters, *d*_*t*_ is the dose rate at time t, *µ* is the half-time for the repair of radiation-induced DNA damage. In fractional radiotherapy, dose is applied as fractions so *d*_*t*_ = d for intervals *t*_2*c−*1_*< t <t*_2*c*_ where c is the fractions per day. Following the studies of Nilsson *et al* and Powathil *et al*, for the low-dose fractionated radiotherapy, Eq (2) can be written as *R*(*n, t*)= *R*_*eff*_ *k*_*R*_*n* where *R*_*eff*_ indicates the effect of c fractions per day and *k*_*R*_ is a factor equals to one when the radiation is applied and otherwise zero^49, 50^.

The effective radiation can be written as:

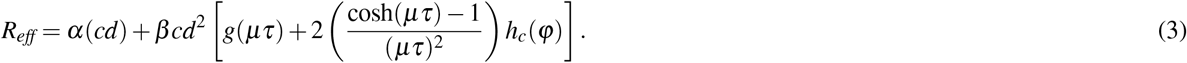

*τ* is the duration of irradiation and the function *φ* = exp(−*µ*(*τ* + Δ*τ*)), where Δ*τ* is the time interval between fractions. The functions *g*(*µτ*) and *h*_*c*_(*φ*) are defined as:

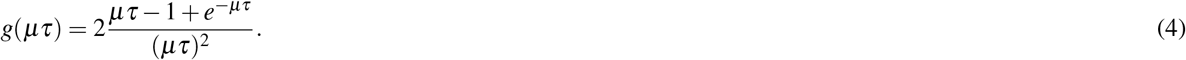

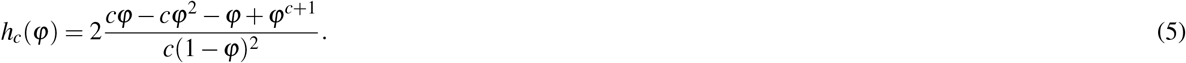

For the initial condition of tumor cells, it is assumed that they are distributed as Gaussian and no-flux boundary conditions are applied.

### Vascular network

Different from normal vessels, the tumor vessel network displays an abnormal structure in terms of function and structure with high permeability and tortuosity. To represent this spatially and temporally heterogeneous structure, a coarse-grained model is used in order to create vessel islands.

In equation (6), the average blood vessel distribution is represented by (m(**x**,t)) and vessel islands are produced as in previous tumor growth models^33, 34, 51^. The second term, *m*(**x**, *t*)[*a*_1_ + *a*_2_*m*(**x**, *t*) + *a*_3_*m*(**x**, *t*)^2^], collapses the equation to the two stable points m=1 and m=0, indicating the areas where vasculature is present and not present. The term 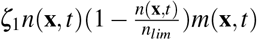 is recruited for generating the sparsity of vessels due to vascular collapse. It is fixed to have nonzero values for the values of *n*(**x**, *t*) greater than *n*_*lim*_. For that range, it attains negative values to eliminate the overgrowth of the vessels, creating a behavior similar to real tumors with high vessel density in tumor periphery and low density in tumor core due to the collapse of vessels in there with the increased solid pressure and interstitial fluid pressure (IFP)^52^.

Angiogenesis is represented with the term of *ζ*_2_∇(*m*.∇*n*) indicating that the vessels move through the interior regions of tumor. The term, *ζ*_3_*m*(**x**, *t*)*A*(**x**, *t*), describes the reaction of tumor vasculature to antiangiogenic agent *A*(**x**, *t*), leading to the elimination of tumor vessels with the application of antiangiogenic agents. The last term accounts for the destruction of vessels due to irradiation, where *ε*_*m*_(**x**, *t*) is a binary map indicating whether vasculature at any point is a tumor vessel or not.

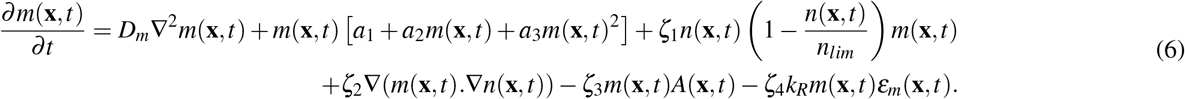

For the initial condition of tumor vessels, it is considered that the vascular network has a randomly positive distribution and no-flux boundary conditions are used for tumor vessels. The related parameters about tumor cell density and vasculature are listed in Table 2. The reaction term, *γ*_1_, between tumor cells and radiotherapy, and the interaction term, *ζ*_4_, between vasculature and radiotherapy are adjusted to observe similar results as in the following study^50^.

**Table 2.** Parameters related to tumor cell and vasculature.

| Parameter | Unit | Value |
| --- | --- | --- |
| $D_n$ | mm<sup>2</sup>/day | $3.5 \times 10^{-2,a}$ |
| $\rho$ | 1/day | $0.35^a$ |
| $n_{lim}$ | 1/mm<sup>2</sup> | $2 \times 10^{6,a}$ |
| $\gamma_2$ | 1/day | $0.19^a$ |
| $D_m$ | mm<sup>2</sup>/day | $1.75 \times 10^{-4,a}$ |
| $a_1$ | 1/day | $-0.35^a$ |
| $a_2$ | 1/day | $1.05^a$ |
| $a_3$ | 1/day | $-0.70^a$ |
| $\zeta_1$ | 1/day | $4.38 \times 10^{-8,b}$ |
| $\zeta_2$ | mm<sup>2</sup>/day | $4.35 \times 10^{-11,b}$ |
| $\zeta_3$ | 1/hours | $1^b$ |
| $m_{lim}$ | - | $\sqrt{2}^a$ |

### Antiangiogenic agent and oxygen concentration

The transport of antiangiogenic agent *A*(**x**, *t*) can be modeled as:

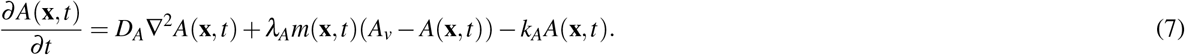

In the equation, the first term is used for the diffusion of antiangiogenic agents from interstitial space, where *D*_*A*_ is the diffusion coefficient of antiangiogenic agents in tissue. The second term represents the transport of the agents from plasma to the tissue, where *λ*_*A*_ and *A*_*v*_ indicate the transvascular diffusion coefficient of antiangiogenic agents and the concentration of antiangiogenic agents in plasma respectively and the last term is used for the natural decay of the agents where *k*_*A*_ is the decay rate of antiangiogenic agents.

The antiangiogenic agents are administered to the plasma with bolus injection by decaying exponentially:

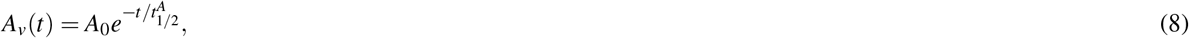

where *A*_0_ is the initial plasma concentration and 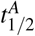 is the plasma half-life of antiangiogenic agent.

A simple reaction diffusion equation is used to depict the spatio-temporal changes in oxygen concentration. In the equation (9), the oxygen concentration is denoted by *K*(*x, t*). The first term on the right hand side displays the diffusion of oxygen through interstitium where *D*_*K*_ is the diffusion coefficient of oxygen. The second term represents the delivery of oxygen by the vasculature leading to the generation of less oxygenated regions (hypoxic) for abnormal tumor vessels that has greater vessel values compared to normal ones. The third term *νK*(**x**, *t*) indicates the uptake of oxygen by tumor cells and the last term is the natural decay of oxygen where *ν* is the decay rate.

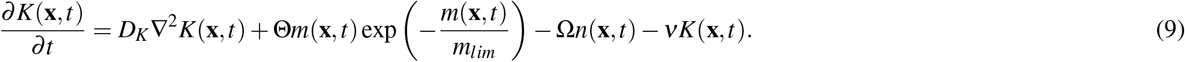

The time scale for the transport of antiangiogenic agents and oxygen is smaller compared to the time scale of the tumor growth, thus antiangiogenic agent and oxygen concentration equations are solved in steady state, i.e. 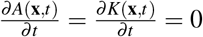. No-flux boundary conditions are applied for both antiangiogenic agent and oxygen equations. Parameters related to radiotherapy, antiangiogenic agents and oxygen are listed in Table 3.

**Table 3.**
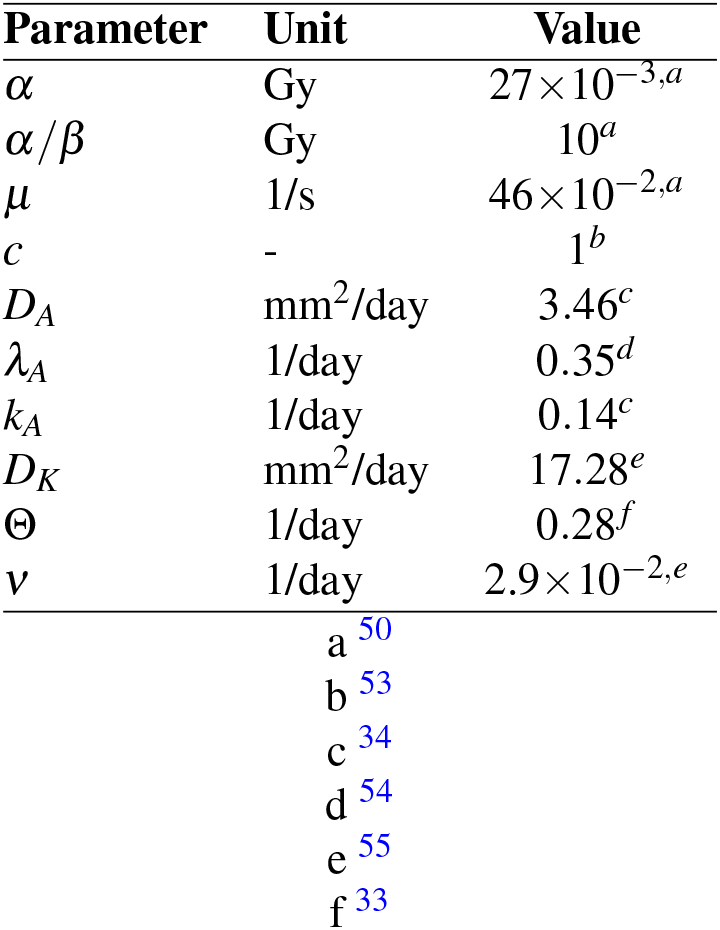
Parameters related to radiotherapeutic values, transport of antiangiogenic agents and oxygen.

## Acknowledgements

This research was supported by TUBITAK grant no. 117F047, Republic of Turkey Ministry of Development grant no. 2009K120520, the Council of Higher Education (100/2000 Ph.D. fellowship), and TUBITAK 2211/A National Ph.D. Scholarship Program.

## Author contributions statement

M.T. and D.Y. collected the data. M.B.U., M.T. and D.Y. analysed the results. All authors reviewed the manuscript.

